# OpenDrive: An Open-Source Microdrive for Large-Scale Neurophysiology

**DOI:** 10.64898/2026.11.09.764949

**Authors:** Gabriel M Stine, Jack Gabel, Nicholas Watters, Mehrdad Jazayeri

## Abstract

Neuropixels-NHP probes have ushered in a new era of high-density neurophysiology recording in large animal models. Here, we present OpenDrive, an open-source microdrive system designed specifically for acute insertion of Neuropixels-NHPs. OpenDrive combines a custom motorized microdrive, six-drive control console, and software platform. Compared with commercially available options, OpenDrive offers (1) low cost, (2) precise, accurate, and stable probe movement, (3) scalability to many-probe recording, and (4) open-source hardware designs and software. In our experiments, OpenDrive has already supported recordings from thousands of neurons across a variety of cortical and subcortical structures. By lowering technical barriers and enabling community-driven development with OpenDrive, we aim to accelerate Neuropixels recording in large animal models.

## INTRODUCTION

Neuropixels-NHP probes are expanding the scale of neurophysiology in large animal models by enabling high-density recordings from multiple brain areas within a single experimental session (Trautmann *etal*., 2025). This technology promises to transform our understanding of the brain. Realizing this opportunity, however, requires hardware that allows laboratories to utilize Neuropixels-NHPs reliably, safely, and at scale.

Adopting Neuropixels-NHPs often requires substantial changes to established experimental workflows. Compared with conventional electrodes, Neuropixels-NHPs have distinct mechanical form factors, mounting requirements, and handling constraints. Their long, thin shanks must be positioned precisely, advanced stably, and protected from lateral forces during insertion. Many existing microdrive systems are poorly suited to these requirements. As a result, laboratories transitioning to Neuropixels-NHPs often need to modify or replace core components of their recording setup, creating a practical bottleneck to adoption.

These challenges motivated us to develop OpenDrive, an open-source microdrive system designed for Neuropixels-NHP recordings. Here, we demonstrate the capabilities of OpenDrive and compare it to two commonly used commercial microdrive systems: the NAN Instruments NAN Drive and the Thomas Recording MEM Drive. We find that OpenDrive offers several advantages over these alternative systems:

1. **Low cost** A complete six-drive OpenDrive system costs approximately $5k to make, whereas comparable NAN Drive and MEM Drive systems cost approximately $50k.
2. **Performance** We show that OpenDrive has similar or improved stability, accuracy, and precision relative to a NAN Drive and a MEM Drive.
3. **Scalability** OpenDrive includes a number of features that facilitate many-probe recording. For example, our software package allows users to (1) create pre-set programs for controlling multiple drives in parallel, and (2) view multiple probe trajectories overlaid on MRI with a real-time display of depth and neural activity (see also Ohayon and Tsao, 2012; Birman *et al*., 2023). Additionally, our system includes a custom probe holder which can accommodate multiple Neuropixels-NHPs on a single microdrive. Using the OpenDrive system, we have been able to lower as many as nine Neuropixels-NHPs into the brain simultaneously.
4. **Open-source** We provide open-source hardware designs and code for all components of the OpenDrive system. To our knowledge, these resources have not been publicly released for any comparable microdrive system.

## SYSTEM OVERVIEW

### Hardware

There are two primary hardware components in the OpenDrive system: the console and the microdrive (Fig. 1). The console receives movement instructions from the graphical user interface (GUI) via USB. It then converts those instructions into motor commands and communicates them to the microdrives through a custom cable. The microdrives then raise or lower their carriages according to the console commands. The microdrives are aligned to their target brain areas through either a custom 3D-printed mount or our optional manual positioning system. Finally, the probes are supported by probe holders, which attach to the microdrive carriages. These hardware components offer multiple beneficial features (Table 1).

**Table 1.**
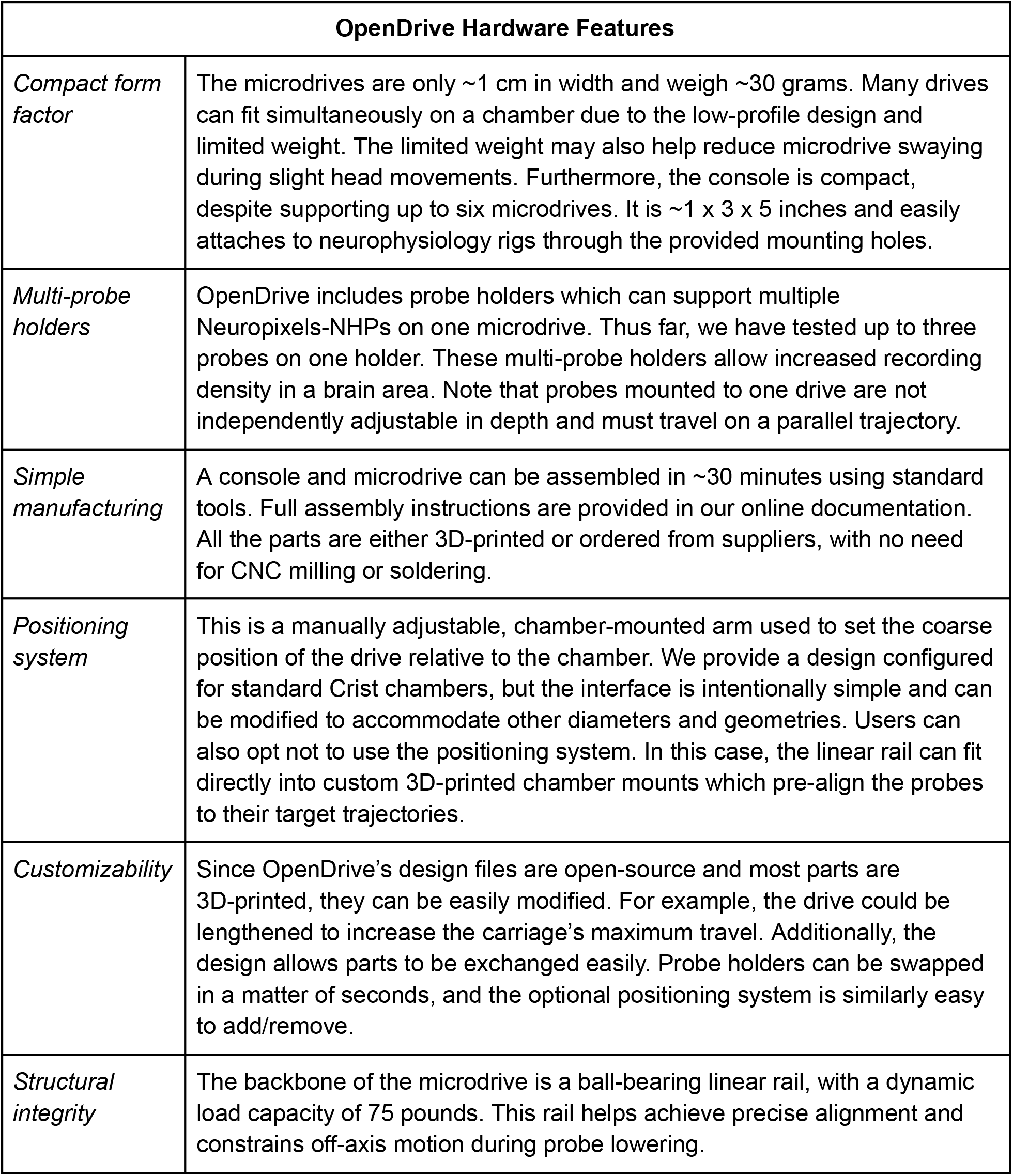
An outline of OpenDrive hardware features.

**Figure 1.**
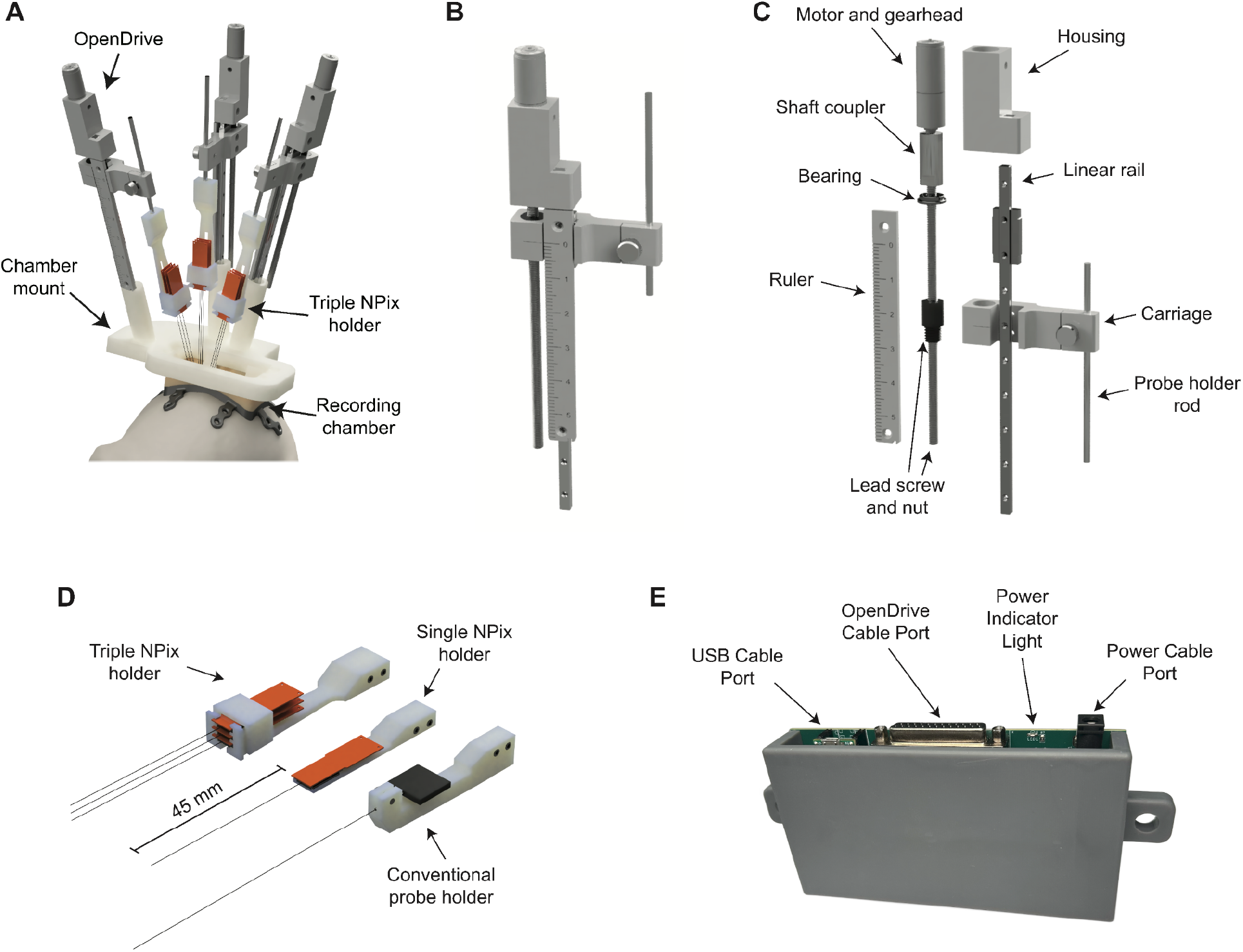
**(A)** A CAD rendering of an OpenDrive system for recording from nine probes simultaneously. Three OpenDrives are held by a 3D-printed mount. The mount is attached to a recording chamber and is designed to pre-align the probes to their target areas in the brain. There are three probes on each drive, which are held by custom probe holders. **(B)** A CAD rendering of a single drive. **(C)** An “exploded” view of B depicting the individual components of the drive. **(D)** Varieties of OpenDrive probe holders. **(E)** Image of the OpenDrive console, which controls up to six drives.

### Software and Firmware

Users control the OpenDrives through a custom Python GUI (Fig. 2). This GUI then sends commands via USB to a microcontroller in the console. The microcontroller runs C++ and handles real-time motor control with low latency. The microcontroller also sends updates back to the GUI to display position information for the user. The OpenDrive GUI and console firmware afford an array of helpful features described in Table 2.

**Table 2.** An outline of OpenDrive software features.

| **OpenDrive Software Features** |  |
|---|---|
| *Trajectory view with SpikeGLX integration* | Users can view their probe trajectories overlaid on an MRI (or any other anatomical scan). A drive’s target depth can be selected directly within the MRI’s coordinate system. Furthermore, multiple trajectories can be viewed on one GUI page to help manage multi-probe sessions. The estimated depth of each probe is updated in real-time and overlaid on the corresponding MRI trajectory. A neural activity heatmap is also overlaid in real-time on the simulated probes for users to check the alignment of neural activity with anatomical landmarks. |
| *Cross-platform compatibility* | The GUI can be run on Windows (which includes SpikeGLX integration) or Mac and does not require a high-end computer because all low-latency operations occur within the console. |
| *Safety* | The GUI and firmware include many safety mechanisms. For example, there are speed and depth limitations. Also, the GUI and console send ‘heartbeat’ messages to one another and will shut off microdrive movement if a message is missed. Finally, all data in the GUI is continuously saved in case the computer crashes. The last-known data will populate when the GUI is reopened. |
| *Remote keypad control* | The GUI offers key bindings. This allows users to control their drives with a wireless keypad while working directly with the subject, away from their OpenDrive computer. |
| *Pre-programmed depth targets* | The GUI includes a mode for executing predefined sequences of positions and speeds. These movement programs can be imported from and exported to CSV files. For example, users can create a program to lower probes into a bath of cleaning detergent and then slowly raise them. |
| *Logging* | Users can log notes about each microdrive within the GUI. These notes, along with all depth information, can be exported as a CSV. |
| *Customizability* | All OpenDrive software and firmware are open-source and can be customized for individual users’ needs. |

**Figure 2.**
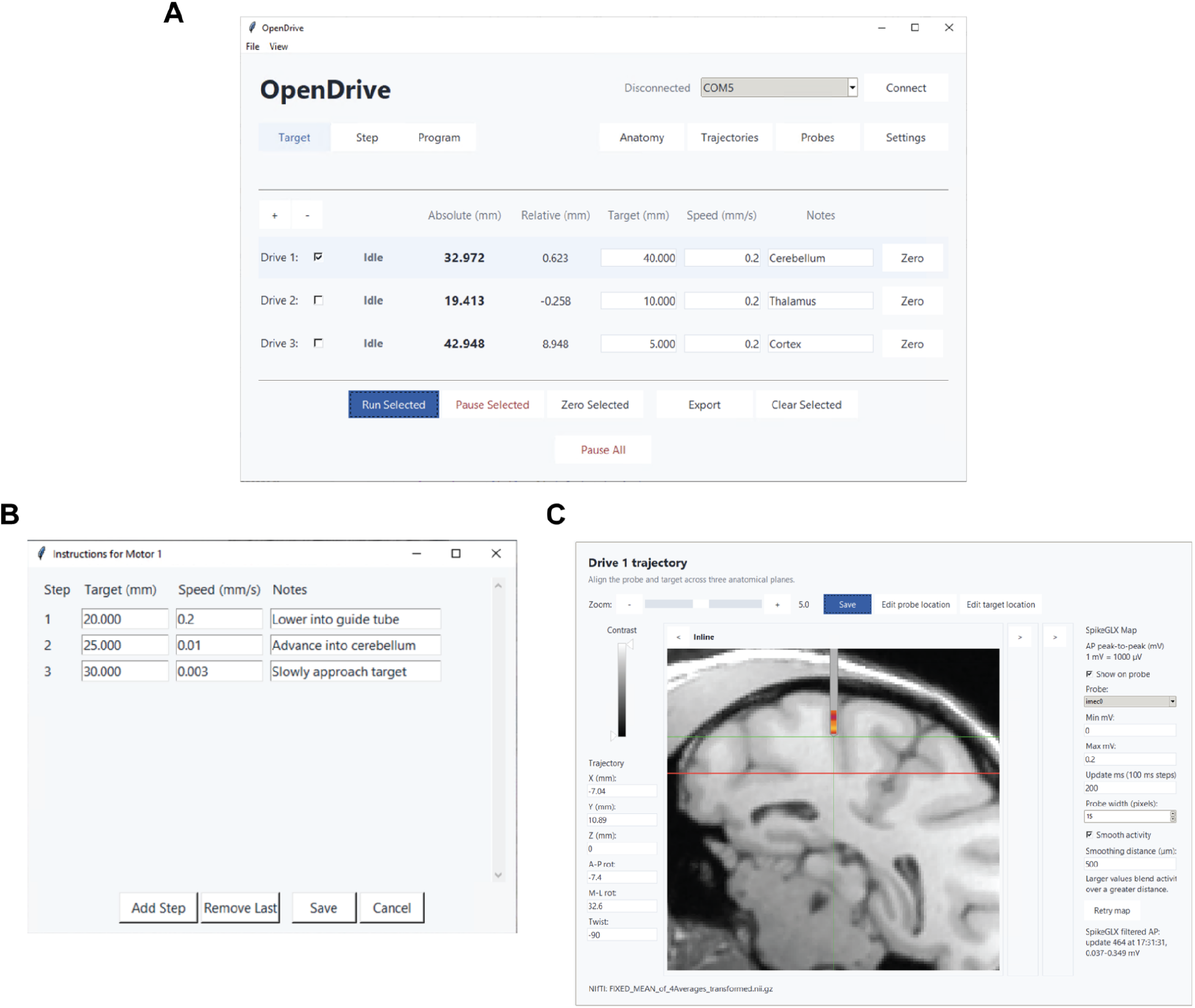
OpenDrive Software GUI. **(A)** Main GUI window set to “Target Mode.” **(B)** Example “instructions” for a drive operating in “Program” mode. **(C)** Screenshot of the “Trajectory Viewer.” OpenDrive integrates with anatomical data and neural data from SpikeGLX. A trajectory for a given drive is defined by the parameters on the left side of the window. The probe position is displayed on the MRI and updated as the drive is running. A real-time neural activity heatmap is overlaid on top of the probe. The drive’s target depth (red line) can be selected and edited within MRI coordinates.

**Figure 2. OpenDrive Software GUI. (A)** Main GUI window set to “Target Mode.” **(B)** Example “instructions” for a drive operating in “Program” mode. **(C)** Screenshot of the “Trajectory Viewer.” OpenDrive integrates with anatomical data and neural data from SpikeGLX. A trajectory for a given drive is defined by the parameters on the left side of the window. The probe position is displayed on the MRI and updated as the drive is running. A real-time neural activity heatmap is overlaid on top of the probe. The drive’s target depth (red line) can be selected and edited within MRI coordinates. These features lower the barriers to conducting large-scale Neuropixels-NHP experiments. To establish whether the OpenDrive system also meets the performance demands of these experiments, we have conducted a series of quantitative tests.

## METHODS

### Lateral motion during lowering

Lateral motion was measured for one OpenDrive, NAN Drive, and MEM Drive using a laser-based assay. A laser was rigidly attached to each drive’s carriage and directed toward a position-sensitive receiver (Thorlabs PDP90A) located at a consistent starting distance from the drive (∼90 mm).

As the drive was lowered, lateral or rotational movement of the carriage displaced the laser spot on the receiver, providing an amplified measurement of deviations from purely axial motion.

Each drive was commanded to travel for ∼2 minutes at 0.005 mm/s, and the two-dimensional position of the laser spot was recorded throughout the movement. Measurements were repeated three times for each drive. To estimate the measurement floor of the apparatus, laser position was also recorded for ∼60 seconds while each drive remained stationary.

Note that we used custom carriage attachments for the NAN Drive and the MEM Drive for all tests to accommodate our measurement instruments.

### Axial positioning accuracy and precision

Axial positioning performance was measured for one OpenDrive, NAN Drive, and MEM Drive. Each drive was mounted rigidly, and displacement along its axis of travel was measured using a plunger-style digital indicator (Mitutoyo ID-C0512ENXB). The plunger was aligned with the direction of drive movement and placed in contact with the moving carriage.

Before each measurement, motor backlash was taken up by approaching the starting position from a consistent direction. Each drive was then commanded to move 10 mm at 0.05 mm/s, and the actual displacement was recorded from the digital indicator. This procedure was repeated 10 times for each drive. Accuracy was quantified as the difference between measured and commanded displacement. All drives undershot the commanded displacement on every trial, so absolute error reflects the magnitude of a consistent directional bias rather than a mix of overshoot and undershoot. Precision was quantified as the standard deviation in measured displacement across repeated movements.

### Use across cortical and subcortical recording targets

We validated OpenDrive in real-world conditions through dozens of recording sessions. Data from one female rhesus monkey (Macaca mulatta; 12 kg) are included here. All experimental procedures conformed to the guidelines of the National Institutes of Health and were approved by the Committee on Animal Care at the Massachusetts Institute of Technology. Three small craniotomies (∼2 mm diameter) were made within a chronically implanted recording chamber to allow access to the cerebellum, thalamus, and parietal cortex. We attached Neuropixels-NHPs to OpenDrives and advanced them into the brain through a stainless-steel guide tube. We used SpikeGLX software (https://billkarsh.github.io/SpikeGLX/) to record Neuropixels data and the drift maps in Figure 4 were computed using MEDiCINe (Watters, Buccino and Jazayeri, 2025) without motion correction.

**Figure 3.**
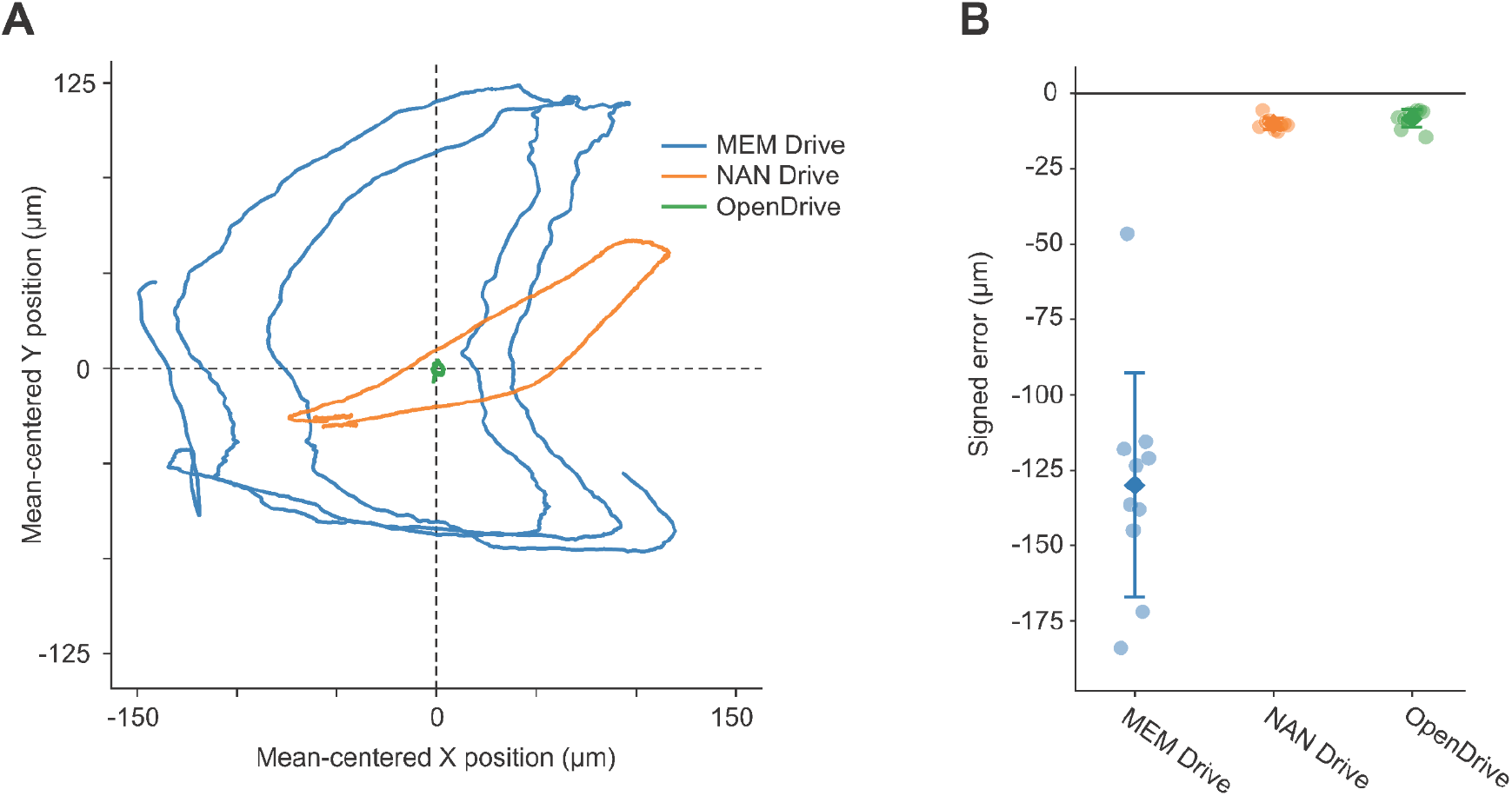
Quantification of microdrive stability, accuracy, and precision. **(A)** Representative mean-centered XY trajectories of the laser spot during one lowering trial for each drive. Traces were smoothed using a 10-sample rolling mean for visualization. The MEM Drive and the NAN Drive produced large looping trajectories, whereas the OpenDrive exhibited minimal lateral displacement of the laser spot. **(B)** Signed positioning error during repeated 10-mm movements of the MEM Drive, the NAN Drive, and the OpenDrive. Points indicate individual trials (n = 10 per drive); diamonds and error bars indicate the mean ± SD. The black line marks zero error. The OpenDrive and the NAN Drive closely approximated the commanded displacement, whereas the MEM Drive substantially undershot the 10-mm command and showed greater trial-to-trial variability.

**Figure 4.**
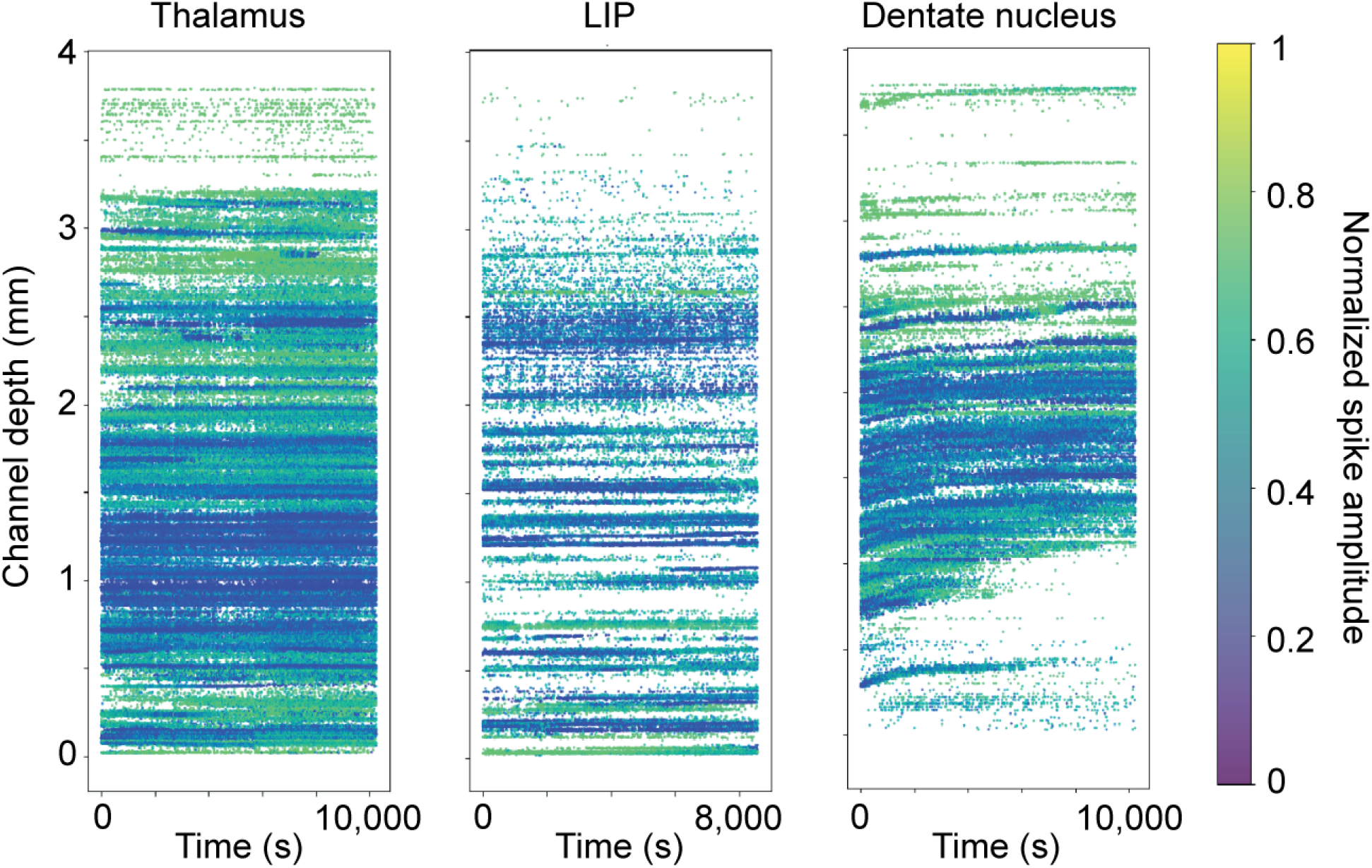
Example drift maps from recordings in the thalamus, lateral intraparietal area (LIP), and dentate nucleus of the cerebellum. Each plot displays normalized spike amplitude as a function of channel depth and time within the session.

## RESULTS

### OpenDrive minimizes lateral motion during lowering

The three drives differed substantially in their lateral motion during lowering (Fig. 3A). Lateral motion was quantified for each trial as the two-dimensional RMS displacement of the laser spot about its mean XY position. Across three trials, RMS laser-spot displacement was 71.65 ± 0.91 µm for the NAN Drive and 95.91 ± 5.36 µm for the MEM Drive. Both the NAN Drive and the MEM Drive produced clear displacement of the laser spot during lowering, with trajectories that followed large, looping paths as the carriage moved along the drive axis. In contrast, the OpenDrive produced minimal displacement of the laser spot during lowering (2.69 ± 0.43 µm). The OpenDrive thus reduced lateral motion by 96% relative to the NAN Drive and 97% relative to the MEM Drive.

In the stationary configuration, RMS laser-spot displacement was low for all three drives (0.72 µm, OpenDrive; 1.59 µm, NAN Drive; 0.71 µm, MEM Drive). We also considered that slight geometric misalignment between the drive axis, laser, and receiver could produce a slow linear shift in the measured laser-spot position during otherwise axial movement. We therefore repeated the analysis after removing trial-wide linear trends separately from X and Y. Linear detrending did not alter the overall ordering of the drives: detrended RMS laser-spot displacement was 2.32 ± 0.23 µm for the OpenDrive, 66.92 ± 3.07 µm for the NAN Drive, and 94.28 ± 4.13 µm for the MEM Drive. Thus, the large differences in lateral motion are not explained by misalignment between the laser, receiver, and drive axis.

These results are consistent with the linear-rail architecture of OpenDrive constraining carriage movement primarily to the intended axis and substantially reducing the large looping trajectories observed in the tested NAN Drive and MEM Drive units.

### OpenDrive achieves accurate and precise axial positioning

The OpenDrive moved accurately and precisely over the tested 10-mm displacement (Fig. 3B). The mean measured displacement for the OpenDrive was 9.992 ± 0.003 mm (mean ± SD), corresponding to an absolute positioning error of 0.008 ± 0.003 mm. Performance was similar for the NAN Drive, which moved 9.990 ± 0.002 mm and had an absolute error of 0.010 ± 0.002 mm. In contrast, the MEM Drive showed greater error and trial-to-trial variability, moving 9.870 ± 0.037 mm with an absolute error of 0.130 ± 0.037 mm.

Absolute positioning error differed significantly among the drives (Welch’s ANOVA, F(2, 15.21) = 51.31, p = 1.73 × 10−^7^). Games–Howell post hoc comparisons found no significant difference between the OpenDrive and the NAN Drive (p = 0.285), whereas both the OpenDrive and the NAN Drive had significantly lower absolute error than the MEM Drive (p = 6.75 × 10−^6^ and p = 7.93 × 10−^6^, respectively). Trial-to-trial variability also differed significantly among the drives (Brown–Forsythe test, F(2, 27) = 7.95, p = 0.0019). Holm-adjusted pairwise comparisons found no significant difference in variability between the OpenDrive and the NAN Drive (p = 0.289), whereas the MEM Drive was significantly more variable than both the OpenDrive and the NAN Drive (adjusted p = 0.030 for both comparisons). Thus, under the tested conditions, the OpenDrive exhibited axial positioning accuracy and precision similar to those of the NAN Drive while outperforming the MEM Drive.

### OpenDrive supports use across cortical and subcortical recording targets

To date, we have successfully used the system to record thousands of neurons across the lateral intraparietal area, dorsomedial frontal cortex, thalamus, cerebellar dentate nucleus, and other regions. These targets span superficial and deep structures and require substantially different probe trajectories and insertion depths, demonstrating that the system can support a range of recording configurations.

For multi-probe experiments, we routinely attach two or three Neuropixels-NHPs to a single drive using the provided multi-probe holder. This configuration allows several probes to be advanced together while limiting the number of drives required around the chamber.

Representative drift maps show qualitatively stable recordings across sessions and brain regions (Fig. 4). In the example sessions, neuronal signals remained at largely consistent estimated depths for ∼3 hours. These results show that OpenDrive has supported repeated, successful recordings from both cortical and deep subcortical targets, including configurations involving multiple drives and multiple probes per drive.

## DISCUSSION

OpenDrive displayed far less lateral motion than both the NAN Drive and MEM Drive, and OpenDrive exhibited axial accuracy and precision similar to or better than the commercial systems. It has also supported routine recordings from thousands of neurons throughout the brain, including lowering up to nine probes simultaneously. Together, these results show that a low-cost, open-source system can provide the performance and scalability required for many-probe Neuropixels-NHP recordings.

OpenDrive builds on several publicly available microdrive systems for neurophysiology in large animal models. Gray *et al*. (2007) developed a computer-controlled, chamber-integrated system for independently advancing multiple conventional electrodes, whereas the PriED system used modular 3D-printed towers to make acute multielectrode recordings less expensive and easier to customize (Patel, Ghose and Eskandar, 2014). More recently, Baeg *et al*. (2021) developed a customizable, MRI-compatible 3D-printed microdrive for neurophysiology, microstimulation, and drug delivery during imaging experiments. These systems demonstrate the value of openly disseminated recording hardware but were designed primarily for conventional electrodes, chamber-integrated arrays, or MRI-compatible applications. OpenDrive was developed specifically for delicate Neuropixels-NHPs and combines a compact motorized microdrive with a six-drive controller and user-friendly software.

The value of an open-source system extends beyond its reduced cost. Electrophysiology hardware is often developed independently within individual laboratories, leading researchers to repeatedly solve similar engineering problems. Open development can instead transform isolated solutions into shared platforms that users can reproduce, adapt, and improve (Siegle *et al*., 2015). Such a culture of open development has indeed been transformative for the rodent neuroscience community (Sanders and Kepecs, 2014; Pachitariu *etal*., 2016; International Brain Laboratory. Electronic address: and International Brain Laboratory, 2017; Mathis *etal*., 2018; Juavinett, Bekheet and Churchland, 2019; Luo *etal*., 2020; Stringer *et al*., 2021; Vöröslakos *etal*., 2021; Madruga *etal*., 2025). This impact motivates our efforts to build a similar culture among groups recording from large animal models, which have relied more on closed-source products. Open development will be particularly consequential for Neuropixels-NHPs, where probe designs, mounting requirements, acquisition software, and experimental configurations continue to evolve. By releasing editable CAD files, PCB designs, firmware, software, assembly instructions, and validation procedures, OpenDrive allows laboratories to adapt the system without waiting for a commercial redesign. Improvements developed by one group, including new probe holders, chamber interfaces, safety features, or software functions, can be tested and returned to the community, reducing duplicated effort and enabling faster iterative development.

OpenDrive nevertheless has limitations. Similar to most commercially available drives, OpenDrive operates without an encoder, so position is estimated from motor commands rather than measured directly, and users must correctly initialize and validate each drive. The system also requires component sourcing, and variation in 3D-printing or assembly could affect performance. Moreover, the present benchmarks examined a limited set of movements on a single example drive of each type. We also did not directly assess tissue displacement, recording yield, or long-term durability, although we have not experienced issues in these domains over months of testing with many OpenDrives. Open-sourcing the system does not by itself resolve these limitations, but it makes them transparent and allows validation, failures, and improvements to be shared across laboratories. We therefore view the present design as a starting point for a low-cost, community-maintained platform that can evolve rapidly alongside large-scale neurophysiology recording tools.

## ACKNOWLEDGMENTS

We thank Aída Piccato, Michael Visconti, Hansem Sohn, and members of the Jazayeri and Shadlen Labs for their help with the design and testing of OpenDrive.

## REFERENCES

Birman, D. et al. (2023) “Pinpoint: trajectory planning for multi-probe electrophysiology and injections in an interactive web-based 3D environment,” eLife, 12, p. RP91662.

Ohayon, S. and Tsao, D.Y. (2012) “MR-guided stereotactic navigation,” Journal of Neuroscience Methods, 204(2), pp. 389–397.

International Brain Laboratory. Electronic address: and International Brain Laboratory (2017) “An international laboratory for systems and computational neuroscience,” Neuron, 96(6), pp. 1213–1218.

Juavinett, A.L., Bekheet, G. and Churchland, A.K. (2019) “Chronically implanted Neuropixels probes enable high-yield recordings in freely moving mice,” eLife, 8, p. e47188.

Luo, T.Z. et al. (2020) “An approach for long-term, multi-probe Neuropixels recordings in unrestrained rats,” eLife, 9(e59716), p. e59716.

Madruga, B.A. et al. (2025) “Open-source, high performance miniature 2-photon microscopy systems for freely behaving animals,” Nature Communications, 16(1), p. 7125.

Mathis, A. et al. (2018) “DeepLabCut: markerless pose estimation of user-defined body parts with deep learning,” Nature Neuroscience, 21(9), pp. 1281–1289.

Pachitariu, M. et al. (2016) “Suite2p: beyond 10,000 neurons with standard two-photon microscopy,” bioRxiv. bioRxiv. Available at: 10.1101/061507.

Patel, S.R., Ghose, K. and Eskandar, E.N. (2014) “An open source 3-d printed modular micro-drive system for acute neurophysiology,” PloS One, 9(4), p. e94262.

Sanders, J.I. and Kepecs, A. (2014) “A low-cost programmable pulse generator for physiology and behavior,” Frontiers in Neuroengineering, 7, p. 43.

Siegle, J.H. et al. (2015) “Neural ensemble communities: open-source approaches to hardware for large-scale electrophysiology,” Current Opinion in Neurobiology, 32, pp. 53–59.

Stringer, C. et al. (2021) “Cellpose: a generalist algorithm for cellular segmentation,” Nature Methods, 18(1), pp. 100–106.

Trautmann, E.M. et al. (2025) “Large-scale high-density brain-wide neural recording in nonhuman primates,” Nature neuroscience, 28(7), pp. 1562–1575.

Vöröslakos, M. et al. (2021) “Metal microdrive and head cap system for silicon probe recovery in freely moving rodent,” eLife, 10(e65859), p. e65859.

Watters, N., Buccino, A. and Jazayeri, M. (2025) “MEDiCINe: Motion correction for neural electrophysiology recordings,” eNeuro, 12(3), p. ENEURO.0529-24.2025.

